# Expansion microscopy for super-resolution imaging of collagen-abundant tissues

**DOI:** 10.1101/2024.02.28.582497

**Authors:** Ya-Han Chuang, Yueh-Feng Wu, Ya-Hui Lin, Yu-Xian Zhou, Shao-Chun Hsu, Sung-Jan Lin, Li-An Chu

## Abstract

Expansion microscopy (ExM) is popular for three-dimensional ultrastructural imaging of cultured cells and tissue slices at nanoscale resolution with conventional microscopes via physical expansion of biological tissues. However, the application of this technology to collagen-abundant thick tissues is challenging. We demonstrate a new method, collagen expansion microscopy (ColExM), optimized for expanding tissues containing more than 70% collagen. ColExM succeeded in 4.5-fold linear expansion with minimal structural distortion of corneal and skin tissues. It was also compatible with immunostaining, allowing super-resolution visualization of three-dimensional neural structures innervating hair follicles and corneas. With ColExM, we succeeded in identifying individual mitochondria and previously unrecognized dendritic spine-like structures of corneal nerves. ColExM also enabled fine mapping of structural rearrangement of tight junctions and actin cytoskeletons. Therefore, this method can facilitate the exploration of three-dimensional nanoscale structures in collagen-rich tissues.

## Introduction

Expansion microscopy (ExM) surpasses the conventional optical resolution limit for imaging biological specimens by expanding them before imaging^1-3^. Through fluorescent labeling, the spatial distribution of molecules of interest can be resolved at super-resolution using conventional fluorescent microscopy. ExM has become increasingly popular for biomedical imaging of cultured cells as well as tissues containing minimal collagen, such as brain slices^1-9^. However, challenges arise when attempting to expand tissues with high levels of collagen, which is a key component of the extracellular matrix in the mesenchyme^1, 3, 10-12^. Collagen content varies among different tissues: less than 5% in the liver, muscles, and kidneys; 10% to 30% in the gastrointestinal tract and aorta; and more than 70% in the skin, cornea, and tendons^13, 14^. Tissues often contain different types of collagen to yield desired mechanical properties. For instance, the skin has an abundance of collagen types I and III, while the cornea predominantly has types I and V^15^. In the original ExM protocol for brain slices, the proteins in the brain samples were first crosslinked to a gel matrix, mainly composed of superabsorbent polymer complexes^4^. The samples were then treated with proteinase K (**ProK**) at room temperature or 37 °C for several hours to one day, resulting in the digestion of long peptides and homogenization of the mechanical characteristics of the specimens. Further incubation in water isotropically expanded the hydrogel and the embedded specimen, increasing the distance between the entrapped molecules^1-3^. Successful expansion of tissues with collagen content ranging from 1% to 5% has been achieved by modified enzymatic digestion of tissue slices^1^. For instance, mouse pancreas, spleen, and lung can be expanded after they are cut into thin slices (< 20 μm thick). Similarly, ∼100-μm thick slices of mouse and human kidney can be expanded by sequential digestion with collagenase and ProK^10^. Thicker ∼250-μm mouse brain slices can be expanded through a two-day ProK digestion treatment^2^. Since only tissue slices can be expanded, the entirety of a three-dimensional tissue sample cannot be imaged. Importantly, expansion of tissues with high collagen content, such as the skin, cornea, pericardium,tendon, and fibrotic tissues, has not been achieved.

In the absence of tissue expansion, other imaging techniques have been employed to capture three-dimensional ultrastructure in tissues with high collagen content^16-18^. Techniques such as focused ion beam and serial block-face scanning electron microscopy have enabled the visualization of three-dimensional ultrastructural features, including neuronal structures and mitochondria in the cornea^16^. However, large-scale ultrastructural mapping, such as visualization of neuronal circuitry, with electron microscopy datasets is limited due to the requirement for tiny sample dimensions. On the other hand, optical super-resolution microscopy offers the resolution previously reserved for electron microscopy, coupled with the advantages of multiplex immunofluorescence staining^17, 18^. However, super-resolution microscopy requires specialized imaging instruments and the use of unique chemical fluorescent dyes. For instance, stimulated emission depletion (STED) microscopy requires a depletion laser^17^, while stochastic optical reconstruction microscopy (STORM) relies on stochastic switching of single-molecule fluorescence signals from a pair of activator–reporter dyes^18^. Regrettably, the imaging depth of super-resolution microscopy is restricted to less than 200 μm due to the limited depth of field of the high-numerical aperture objectives required for this technique^19^. Therefore, the development of an expansion method for collagen-abundant tissues is an excellent solution that is urgently needed.

In this study, we successfully extended the application of ExM to tissues with a collagen content greater than 70%—corneal and skin tissues—which enabled optical super-resolution imaging. Our method, collagen expansion microscopy (ColExM), achieved 4.5-fold linear tissue expansion with minimized structural distortion and was compatible with immunostaining procedures. An additional advantage of ColExM was the highly enhanced tissue transparency, which dramatically improved light penetration into the skin. This was previously challenging with traditional tissue-clearing methods. Furthermore, we were able to successfully reconstruct neural nanostructures within these tissues and map the structural changes in tight junctions in response to calcium deprivation. These advancements open new avenues for detailed exploration of nanoscale structures in collagen-rich tissues.

## Results

### Collagen expansion microscopy allows the expansion of collagen-abundant tissues with minimal tissue distortion

In our investigation, we aimed to determine whether previously reported protocols, designed for expanding tissues containing less than 5% collagen, could be applied to tissues with more than 70% collagen for imaging purposes. Following the method described by Asano et al.^2^, we performed experiments on PFA-fixed corneal tissues. However, despite subjecting the tissues to prolonged ProK digestion at 37 °C for two days, the expansion we observed was nonuniform. While the non-mesenchymal layers, such as the epithelium and endothelium, showed expansion, the corneal stroma remained nearly unchanged in size. Similar inhomogeneity in tissue expansion was also observed in the skin sample **(Supplementary Fig. 1a)**

We then attempted the method proposed by Chozinski et al.^10^, involving ProK digestion at 37 °C for one day followed by collagenase treatment at 37 °C for an additional day. However, similar to the Asano method, this approach also did not result in uniform corneal expansion and the cornea exhibited distortion **(Supplementary Fig. 1b)**. Moreover, considering that collagenase is typically applied before tissue fixation to dissociate tissues into single cells, we also experimented with a 15-minute pre-fixed collagenase treatment. However, this approach also proved ineffective in achieving uniform tissue expansion **(Supplementary Fig. 2)**. These results indicate that current modified expansion protocols, which work well for thinly sliced tissues with a collagen content of less than 5%, are inadequate for achieving uniform expansion of thick tissues with more than 70% collagen content.

Another potential strategy involves modifying the operating temperature of ProK. Most ExM studies traditionally expose tissue to ProK at room temperature or 37 °C for digestion^1-3^. This cautious approach is often taken due to concerns that higher temperatures might excessively disrupt protein epitopes during digestion, potentially compromising further immunolabeling. However, it is worth noting that the activity of ProK is known to increase with temperature, and its optimal working range falls between 50–55 °C^20^. To explore this, we varied the working temperatures and durations of ProK treatment of collagen-abundant tissues. Surprisingly, we found that these tissues could be uniformly expanded under treatment with type 2 collagenase for 48 hours, followed by ProK digestion at temperatures above 50 °C for 24 hours **(Supplementary Fig. 3)**. Using nuclei as an indicator, we observed that the average longest axis of corneal endothelial cell nuclei increased from ∼20 μm before expansion to ∼75 μm after expansion, resulting in an approximately ∼4-fold expansion factor, which was slightly less than the theoretical expansion factor of 4.5 for acrylamide-sodium acrylate expansion gels^4^. We extended the ProK digestion duration to 48 hours and successfully achieved a 4.5× expansion of all the collagen-abundant tissues tested **(Fig. 1a–c)**. Our results demonstrate that by modifying the working temperature and treatment duration of ProK, it is possible to achieve complete expansion of collagen-abundant tissues while preserving the overall tissue structures. Therefore, we used the collagenase treatment for 48 hours at 37 °C and the ProK treatment for 48 hours at 55 °C for subsequent experiments.

**Fig. 1.**
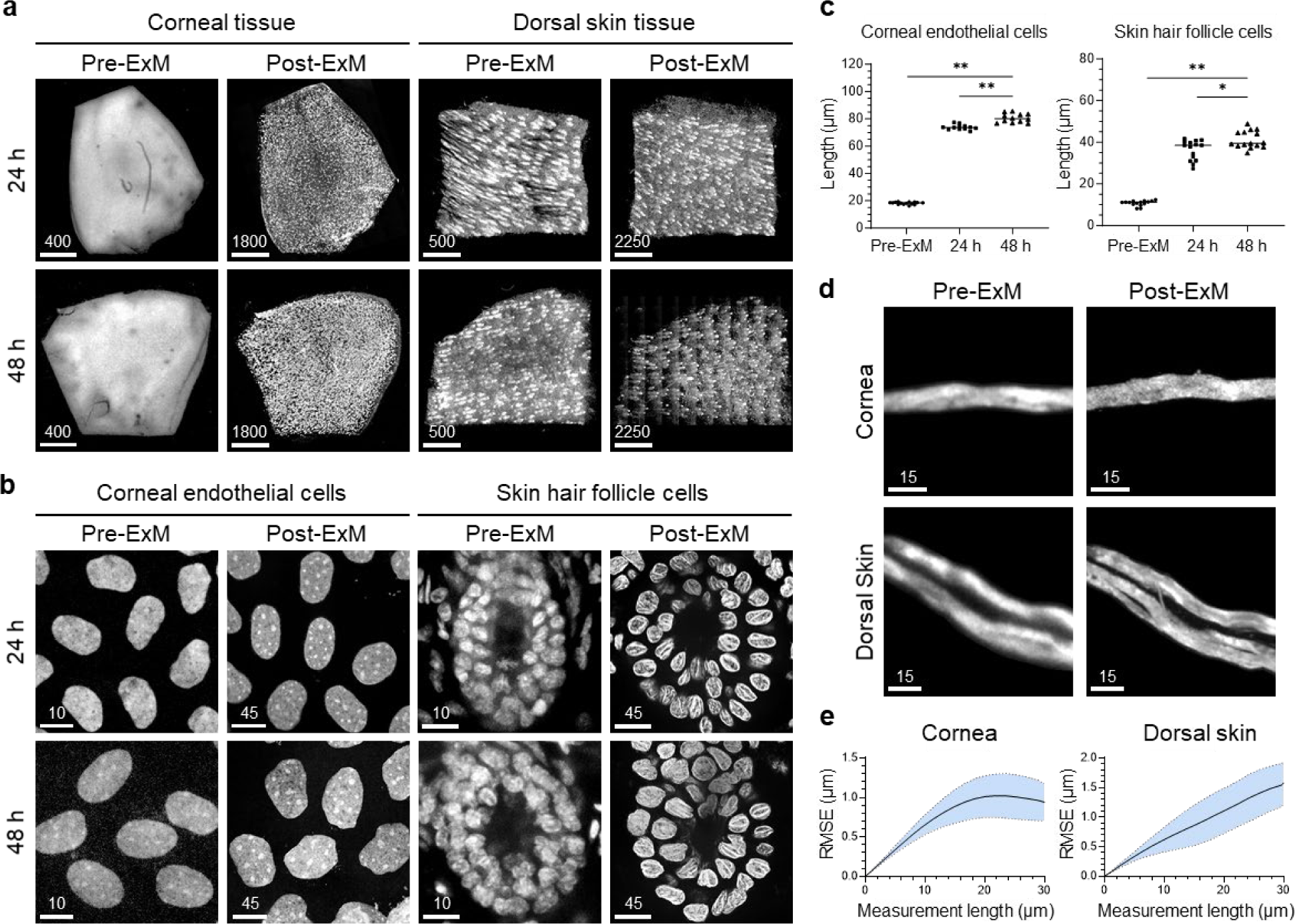
Collagen expansion microscopy (ColExM) of collagen-abundant tissue expansion. **a and b,** Representative images of pre- and post-expansion of corneal and dorsal skin tissues under 24-hour and 48-hour ProK digestion treatments. Nuclei were stained with PI staining and are shown in gray. **c,** Measurement of the longest axis of the cell nucleus to quantify expansion factors of corneal endothelial cells and skin hair follicle cells. We show the mean values from three different experiments; we measured the nuclei of at least 15 cells in each experiment to calculate the corresponding expansion factor. Error bar = ± S.E.M. * p < 0.01, ** p < 0.0001, Student’s *t*-test. Scale bar unit: μm. Scale bars are not adjusted for tissue expansion. **d,** Pre-expansion microscopy (ExM) Airyscan image and post-ExM confocal image of Thy1-positive nerves of the same region in corneal and dorsal skin tissues. **e,** Root-mean-square error analysis of ColExM and Airyscan images of Thy1-positive nerves in the corneal and dorsal skin tissues (blue line, mean; shaded area, standard deviation; n = 3 samples). Scale bar unit: μm. Scale bars correspond to pre-expansion dimensions.

To assess the degree of structural distortion, we performed root-mean-square error (RMSE) analysis of Airyscan confocal images of Thy1-positive nerves in pre- and post-expanded tissues **(Fig. 1d)**. RMSE is a method of quantifying discrepancies between predicted values generated by an estimator and the actual values. Consequently, it has been used to evaluate the extent of tissue distortion before and after expansion^3^. Our data indicated distortion values in the expanded cornea of approximately 6– 7% within 10 μm (**Fig. 1e,** n = 3) and about 3–6% between 10 μm and 30 μm. In contrast, the analysis of dorsal skin revealed distortions ranging from 5 to 7% (**Fig. 1e,** n = 3). A notable “side effect” of expansion microscopy is its ability to enhance tissue transparency by pulling the molecules away from each other. While the traditional tissue-clearing technique can render mouse cornea fully transparent **(Fig. 2a**), it is difficult for it to clear compact structures like mouse skin (**Fig. 2b**). In unexpanded skin embedded in clearing buffer, the signal starts to blur beyond 100–150 μm even after 24 hours. The dense and tightly packed fibrillar structures, such as collagen fibers, can limit penetration by clearing agents and hinder their uniform distribution within tissue. In contrast, after expansion, the skin achieves a high level of transparency, while the relative imaging resolution is increased, thereby enabling intact three-dimensional analysis of the skin structure (**Fig. 2c**).

**Fig. 2.**
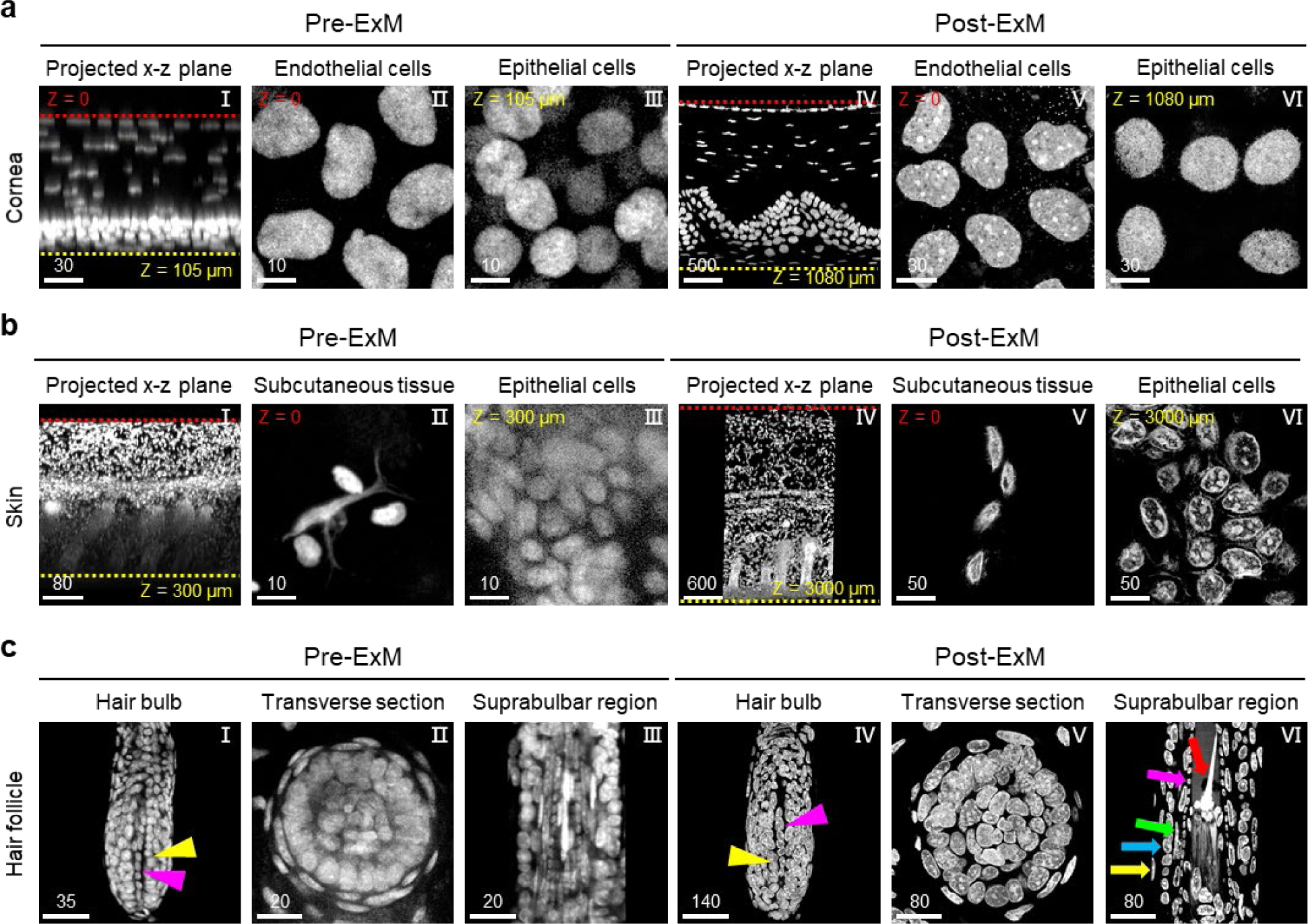
Collagen expansion microscopy (ColExM) enhances tissue transparency. **a and b,** Representative pre- and post-expansion images of the cornea and skin. Images are of the X–Z view and X–Y view of PI-stained tissues. Unexpanded tissues were incubated in NFC-2 solution for 24 hours before imaging. Red dotted lines: the position of Z = 0; yellow and red dotted lines indicating the z depth. **c,** Representative pre- and post-expansion images of the hair follicle in skin. Magenta arrowheads: dermal papilla; yellow arrowheads: germinative cells; red arrow: hair shaft; magenta arrow: inner root sheath cells; green arrow: companion layer cells; blue arrow: outer root sheath cells; and yellow arrow: dermal sheath cells. Nuclear PI staining is shown in gray. Scale bar unit: μm. Scale bars are not adjusted for the expansion factor.

By exploiting the benefits of tissue transparency and increased relative imaging resolution of skin, we further extended the application of ColExM to investigate the intricate cellular arrangement within anagen hair follicles. The epithelium of anagen hair follicles forms a cylinder with eight concentric layers^21^; starting from the periphery, these include the dermal sheath, outer root sheath, companion layer, inner root sheath, and the hair shaft. The hair bulb, in the proximal portion of the anagen hair follicle, comprises mesenchymal cells, referred to as the dermal papilla, which are encompassed by germinative cells. However, a clear visualization of the entire 3D structure of hair follicles remains challenging due to their tightly packed multi-layered nature. To avoid melanin interference, we imaged the hair follicles from the dermal layer to the epidermal layer. However, we could only visualize the hair bulb, including the dermal papilla (magenta arrowheads) and germinative cells (yellow arrowheads), before expansion **(Fig. 2c I),** but not the detailed structure of the proximal part of the hair shaft. In contrast, the detailed structure and cell composition of the proximal region of the hair shaft became clear in the post-expanded sample (red arrow) **(Fig. 2c VI)**. In addition, the morphology of cell nuclei in the cross-section of post-expansion hair follicles became sharper than that in pre-expansion hair follicles **(Fig. 2c II)**. In the post-expansion hair follicles, the horizontal inner root sheath cells (magenta arrow) enwrapping the hair shafts could also be visualized, while companion layer cells were flattened vertically (green arrow) **(Fig. 2c V)**. Compared to the inner root sheath and companion layer, outer root sheath cells appeared rounder and larger (blue arrow) **(Fig. 2c VI)**. The dermal sheath cells were distributed discontinuously on the most peripheral side (yellow arrow) **(Fig. 2c III)**. The gaps between dermal sheath cells were often less than 100 nm; therefore, they could be clearly demarcated only after expansion. In summary, ColExM enabled the expansion of skin, cornea, and hair follicles while facilitating the visualization of different cell types and tissue structures in detail.

### Visualization of nerve nanostructure in epithelial tissues by collagen expansion microscopy

We used ColExM to investigate whether the fluorescent protein signal and immunostained dye signal remain preserved after tissue expansion. We first directly used ColExM with Thy1-YFP-H transgenic mice, in which YFP is highly expressed in motor and sensory neurons, along with subsets of central neurons. However, the YFP signal quenched dramatically even with the protein retention anchor **(Fig. 3a)**, consistent with Tillberg and colleagues’ findings^22^. Therefore, we employed a streptavidin-based amplification technique to enhance the YFP fluorescent signals. In post-expansion samples of the cornea and skin with the boosted YFP signal, we were able to observe multiple layers of neurons wrapping around cells, a feature that was indistinguishable in the pre-expansion tissues **(Fig. 3b I, V and Supplementary video 1)**.

**Fig. 3.**
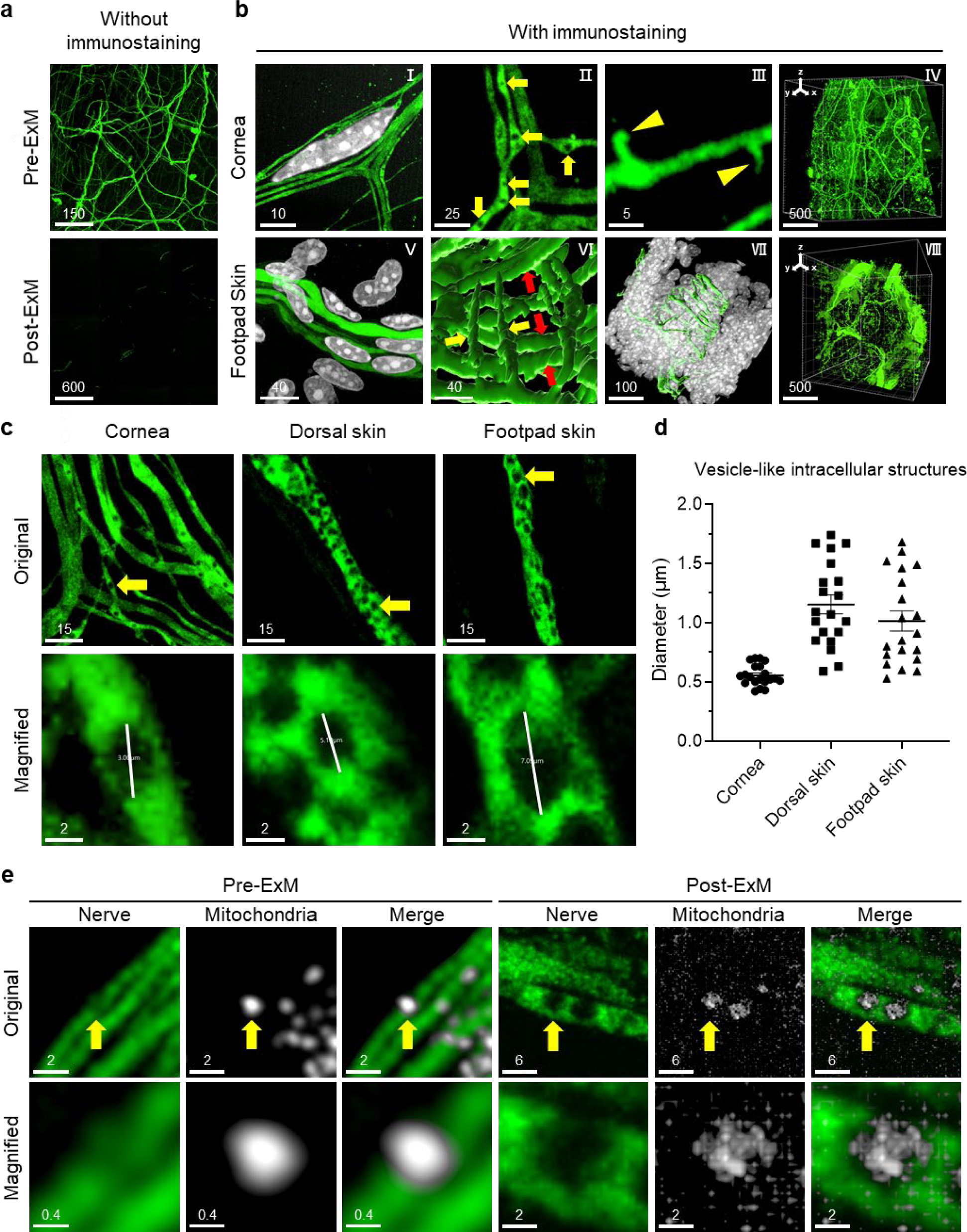
Visualization of mitochondria in Thy1-positive nerves in collagen-abundant tissues after expansion. **a,** Pre-expansion microscopy (ExM) and Post-ExM images of Thy1-positive nerves in the cornea without immunostaining. **b,** Post-ExM images of YFP-labeled nerves in the cornea and footpad skin of Thy1-YFP-H transgenic mice. Nuclear DAPI staining is shown in gray. Nerve bundles (I, V); vesicle-like intracellular structures in corneal nerves (yellow arrows) (II); spine-like structures (yellow arrowheads) of corneal nerves (III); Circumferential (red arrows) and longitudinal (yellow arrows) YFP-labeled nerves (VI); YFP-labeled nerves surrounding the hair follicle in skin (VII); and 3D visualization of YFP-labeled nerves (IV, VII). **c,** Post-ExM images of vesicle-like intracellular structures in corneal and skin nerves. **d,** Measurement of the diameter of vesicle-like intracellular structures in Fig. 3c correspond to pre-expansion dimensions. Error bar = ± S.E.M. **e,** Confocal images of tubulin β3-labeled nerves (green) and GFP-labeled mitochondria (gray) in the cornea before and after expansion. Scale bar unit: μm. Scale bars are not adjusted for the expansion factor.

The sensory nerve innervates the corneal stroma and corneal epithelium^23^. Through ColExM, we observed multiple layers of Thy1-positive nerve fibers enwrapping the nuclei of fibroblast-like cells **(Fig. 3b I and Supplementary video 1)**. These nerve fibers contained dendritic spine-like structures (yellow arrowheads) **(Fig. 3b III, Supplementary video 2)**, a novel observation that has not been reported before. Consistent with previous research^24^, we identified two types of Thy1-positive sensory nerves surrounding the hair follicles based on their fiber orientation: circumferential (red arrows) and vertical (yellow arrows) in the outer and inner layers, respectively **(Fig. 3b VI, VII and Supplementary videos 3 and 4)**. Interestingly, we discovered through ColExM several cell nuclei sandwiched between circumferential and longitudinal YFP-labeled nerves **(Supplementary video 5)**. In addition, the nerves innervating the sweat glands in the footpad were also clearly visualized **(Supplementary video 6)**.

We also discovered rhythmic vesicle-like intracellular structures deficient in fluorescent signals within these neurons **(Fig. 3b II and 3c)**. These intracellular structures in the expanded cornea were highly similar in size and shape, with an average diameter of ∼0.5μm (2.5 μm post-expansion). Notably, they were approximately equidistant from each other. In contrast, the distribution, size, and shape of these vesicle-like intracellular structures in the expanded dorsal and footpad skin differed considerably, with irregular morphology and diameter ranging from ∼1 to 1.2 μm (4.5 to 5 μm post-expansion) **(Fig. 3d)**. Next, we investigated whether the vesicle-like intracellular structures contained mitochondria. We stained corneas from mito-GFP mice^25^, a mitochondrial reporter of cytochrome c oxidase 8A (COX8A) fused with GFP, with tubulin β3 antibody to label the neurons. COX8A is expressed within mitochondrial complex IV^26^. Before expansion, the vesicle-like intracellular structures in the corneas of mito-GFP mice stained with tubulin β3 were not distinctly discernible; nonetheless, some mito-GFP signals were observed in proximity to the nerves (yellow arrows). After expansion, both the mito-GFP and nerve signals became more distinct. Moreover, the mito-GFP signals populated the vesicle space (yellow arrows), suggesting that the vesicle space in the nerve contained mitochondria **(Fig. 3e)**. In summary, ColExM enabled the visualization of nanoscale nerves innervating hair follicles and the cornea, as well as dendritic spines and mitochondria in corneal nerves.

### Three-dimensional ultrastructural visualization of tight junctions in the corneal endothelium by collagen expansion microscopy

Tight junctions are located along the cell membrane in the uppermost region of the lateral plasma membrane to connect neighboring cells and control the passage of molecules and ions through the paracellular space^27^. Dependent on homophilic and heterophilic composition, transmembrane proteins, consisting of claudins, occludins, and junctional adhesion molecules, create cation, anion, or water-selective pores. Intracellular proteins, the zonula occludens (Zo) complex, are present as homodimers or Zo1/Zo2 and Zo1/Zo3 heterodimers connecting to transmembrane proteins^28^. The N-terminus of the Zo complex directly binds to the actin cytoskeleton^29^. Instead of the entire tight junction structure, only tight junction strands formed by transmembrane proteins in various epithelial and endothelial tissues, such as the intestinal epithelium^30^, corneal epithelium^31^, skin epithelium^32^, and the corneal endothelium,^33^ can be examined by transmission electron microscopy (TEM). These structures reveal electron-dense plaques on the cytoplasmic side of tight junctions, consisting of the Zo complex and the actin cytoskeleton. After tissue expansion by ColExM, the overall hexagonal distribution of Zo1 remained intact, and the fine structure exhibited a delicate zigzag line instead of a straight line as seen in the unexpanded sample. This observation is consistent with the regularly spaced distribution of tight junctions along the cell membranes revealed by TEM^34^. In addition, F-actin displayed a more filamentous appearance and was located on the side of Zo1 **(Supplementary video 6),** rather than the co-localized pattern observed in the pre-ExM images **(Fig. 4)**.

**Fig. 4.**
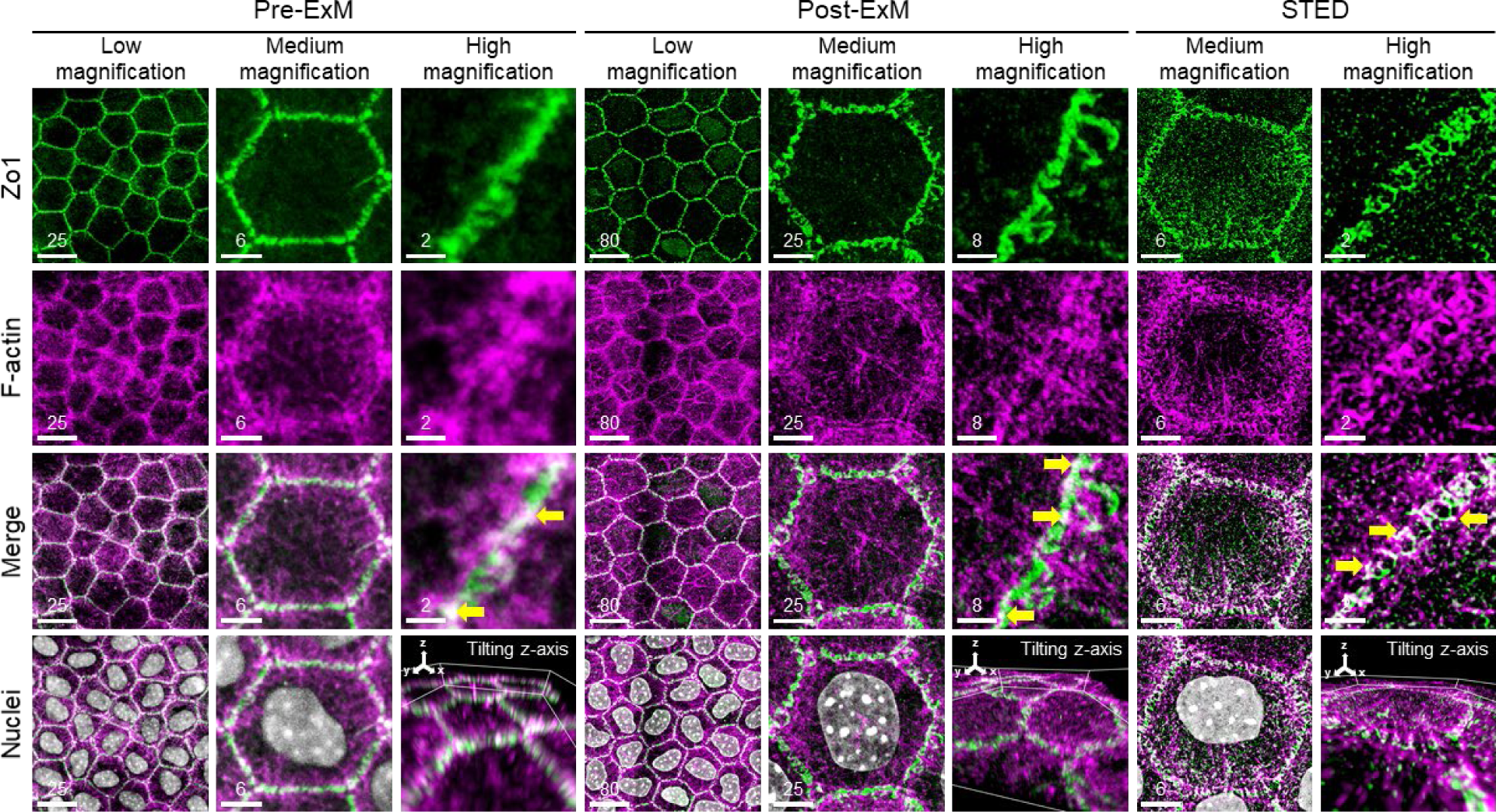
Confocal images of tight junctions in the corneal endothelium of expanded mouse corneas. Confocal images of Zo1 (green), F-actin (magenta), and nuclei (gray) in the corneal endothelium before and after expansion and STED microscopy images of unexpanded corneas. Colocalization of Zo1 and F-actin is shown in white in merged images (yellow arrows). Scale bar unit: μm. Scale bars are not adjusted for the expansion factor.

We then compared the images obtained with ColExM and those obtained with super-resolution microscopy. In the case of super-resolution microscopy, we used STED microscopy with high-numerical aperture (1.4 NA) and high-magnification (100×) objectives to observe immunostained unexpanded corneas. When we employed a doughnut-shaped focal pattern generated by the depletion laser, Zo1 signals in the STED images also exhibited a zigzag pattern **(Fig. 4)**. However, compared to ColExM images, the background noise in STED images was significantly higher. Zo1 signals not only appeared at the cell border but were also detected dispersed in the cytoplasm. Moreover, the F-actins labeled with phalloidin in the STED images were more heavily co-localized with Zo1 signals than those in ColExM images. The STED images revealed a highly similar pattern for Zo1 and F-actin at the cell border, perpendicular to the cell membrane. This result contradicts the findings of a previous study, which showed F-actin linking only to 28 amino acids of the Zo1 protein in the cytoplasm near the cell border^29^. On the other hand, in ColExM images, F-actin exhibited a trail-like structure along the cell membrane border, in a parallel alignment to the Zo-1 signal. In addition, inside the cell, F-actin adopted a radial shape, perpendicular to the F-actin observed on the membrane, suggesting an inward force exerted by the F-actin to maintain the cell’s shape. These findings demonstrate that ColExM can achieve a resolution similar to that of STED while preserving a superior molecular distribution and signal-to-noise ratio in microscopy images.

### Disruption of tight junctions in the cytoplasm of the corneal endothelium following calcium deprivation

Calcium ions are essential for the barrier functions of tight junctions^35^. Depletion of calcium ions by treatment with the calcium chelator ethylene glycol tetraacetic acid (EGTA) reduces transepithelial/transendothelial electrical resistance (TEER) and increases paracellular/transcellular permeability^36, 37^. EGTA disrupts tight junction transmembrane proteins by reducing the affinity of claudins in heterophilic forms^38^, including occludin dephosphorylation^39, 40^ and causing occludin/Zo1 redistribution in cultured cells^41, 42^. However, the way in which the intracellular Zo1 and actin cytoskeleton in the corneal endothelium respond to calcium depletion remains unclear. Therefore, we used ColExM to map the process of tight junction disruption at a super-resolution level. Mouse corneas were bathed in a culture medium containing EGTA for different durations before further processing for imaging.

Without expansion, the width of the Zo1 belt between cells increased after 30 minutes of EGTA treatment, as observed by conventional confocal microscopy. Similarly, the F-actin signals gradually became blurred after the 30-minute EGTA treatment. Both F-actin and Zo1 formed a thicker hexagonal belt at the cell border, with the co-localized proportion gradually decreasing during the EGTA treatment process. However, due to resolution limitations, we could only confirm that these two proteins dispersed within the cytoplasm. The exact cause of Zo1 diffusion within the cytoplasm, possibly due to F-actin tension, could not be clearly demonstrated in this set of images **(Fig. 5a)**.

**Fig. 5.**
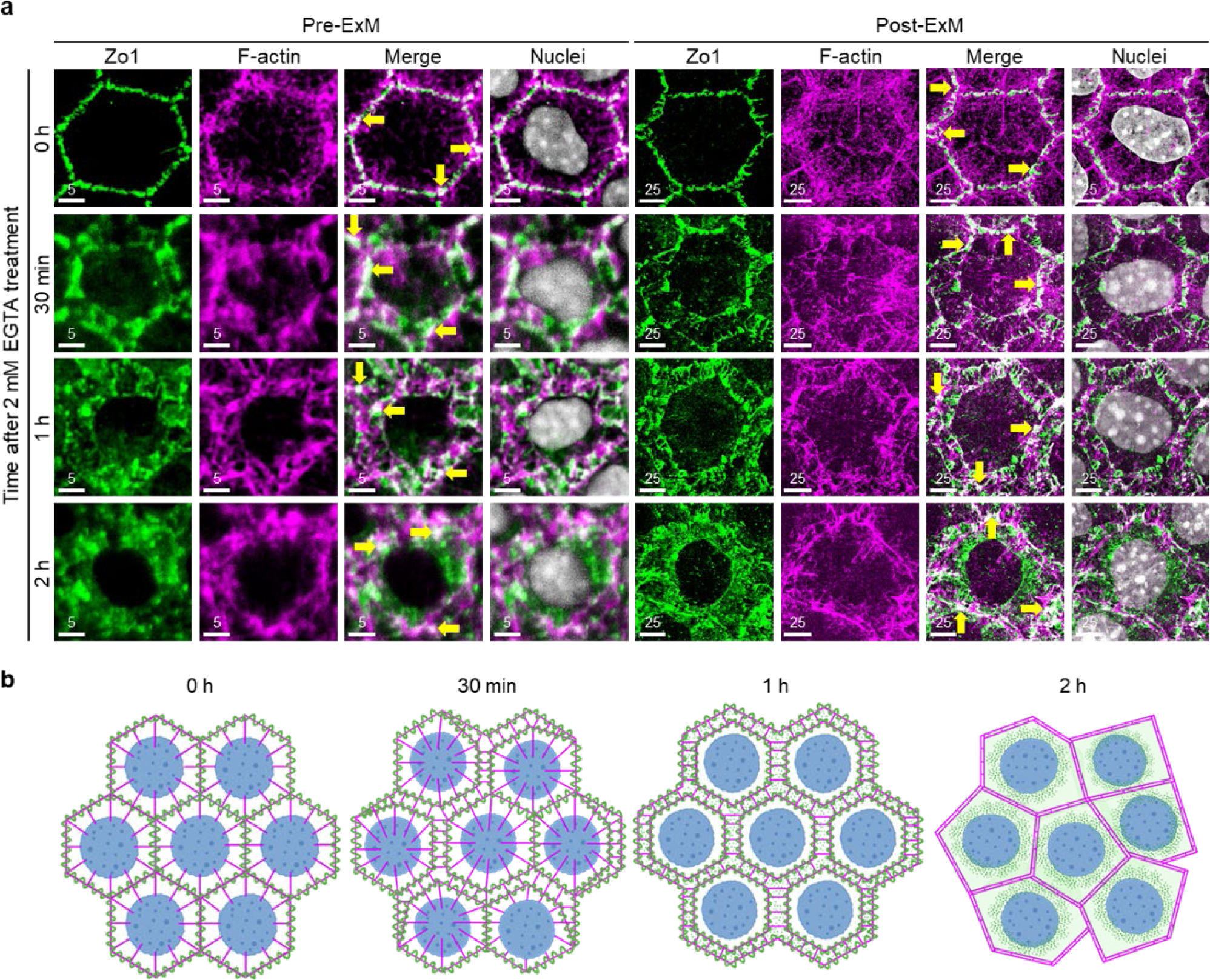
Tight junctions of corneal endothelial cells conformationally change after disruption by EGTA. **a,** Expression of Zo1 (green), F-actin (magenta), and nuclei (gray) in the corneal endothelium before and after expansion without EGTA treatment and following EGTA treatment for 30 minutes, 1 hour, and 2 hours. Colocalization of Zo1 and F-actin is shown in white in merged images (yellow arrows). Scale bar unit: μm. Scale bars are not adjusted for the expansion factor. **b,** Schematic diagram of the tight junction conformation change process. Zo1 is indicated in green, F-actin is indicated in magenta, and the cell nucleus is indicated in blue.

Images obtained from post-ColExM samples revealed a distinct F-actin pattern in both untreated groups and those treated with EGTA for 30 minutes. This pattern displayed a robust radial filamentous arrangement, connected to the F-actin at the cell border and centered around the cell nucleus. This particular configuration was unidentifiable in images acquired prior to expansion. Conversely, due to the ineffectiveness of EGTA in dephosphorylating intercellular occludin, a noticeable change occurred after the 30-minute EGTA treatment. Zo1 proteins from neighboring cells started to segregate into two to three hexagonal segments on each cell. This phenomenon might be attributed to the ongoing force exerted by intracellular F-actin on Zo1, as evidenced by the partial co-localization of Zo1 and F-actin along the cell border during this period. Following exposure to EGTA for one hour, Zo1 exhibited a marked separation from neighboring cells along all six hexagonal sides, and the previously observed radial arrangement of F-actin within the cell ceased. Following EGTA treatment for two hours, Zo1 showed uniform dispersion throughout the cytoplasm. Meanwhile, F-actin underwent rearrangement, assuming a hexagonal configuration along the cell border, and the formerly evident radial filamentous pattern was no longer discernible, indicating the loss of the force binding F-actin to the Zo1 protein during this particular stage (**Fig. 5a**). These data revealed that tight junction disruption resulted in the deposition of Zo1 in the cytoplasm, and the F-actin pulling force that initially maintained the Zo1 structure was disrupted during EGTA treatment of occludin. ColExM enabled the mapping of the entire process of Zo1 and F-actin in the corneal endothelium after EGTA treatment **(Fig. 5b)**.

## Discussion

Collagen expansion microscopy enabled 4.5× expansion of collagen-abundant tissues like the mouse cornea and skin (dorsal/footpad). This expansion was successfully achieved by optimizing the working temperature and duration of ProK and collagenase digestion. We demonstrated that ColExM improved resolution while preserving molecular distribution, outperforming techniques such as STED^17^ and STORM^18^. This capability fulfills the demand for 3D ultrastructure visualization unmet by techniques like TEM or FIB in this field. 3D images of various neuronal ultrastructures embedded in collagen-abundant tissues can be visualized distinctly, enabling further quantitative analysis. Furthermore, we have broadened the range of applications of ColExM to characterize not only delicate neuronal structures but also the conformational changes in the intracellular domain within the corneal endothelium following tight junction disruption by EGTA treatment.

To achieve nanoscale multiplex staining of tissues, optical super-resolution imaging serves as an alternative solution that eliminates the need for an expansion process. However, our investigations indicated that this approach may not be suitable for studying weak signals. When we used STED, an optical super-resolution method, to visualize the interaction between Zo1 and F-actin, we observed that while the Zo1 channel exhibited a strong signal, the F-actin signals appeared scattered and discontinuous, posing challenges to the differentiation between the actual signal and background noise **(Fig. 4)**. Although the scattering of the F-actin signal could be attributed to the sparse labeling of phalloidin to F-actin^43^, the integration of the depletion laser in STED amplified both the discontinuous F-actin signals and the background noise. These findings suggest that current super-resolution methods may encounter significant challenges when dealing with weak original fluorescent signals. In contrast, ExM tends to avoid such difficulties. In ColExM, we employed a streptavidin-based technique to enhance fluorescent signals, thereby mitigating the reduction in fluorescent signals caused by increased molecular distances during the expansion process. Due to its compatibility with immunostaining, ColExM is also possibly compatible with commercially available organelle-staining dyes, such as Lysotracker^44^ and MitoView^TM^ Green^45^. For example, the incorporation of MitoView^TM^ Green into ColExM of clinical specimens may offer the potential to explore the mechanisms of mitochondrial dysfunction and mitophagy in the pathogenesis of Fuchs endothelial corneal dystrophy in future studies.

In initial observations of ultrathin sections using TEM, tight junctions appear as condensed signals at zones where plasma membranes of neighboring cells focally make complete contact^46^. Examination of the connection between Zo1 and F-actin using TEM remains a challenge due to the characteristic coil structure and restricted dimensions of ultrathin sections. However, the colocalization of Zo1 and F-actin has been visualized in lipid bilayers of giant unilamellar vesicles using total internal reflection fluorescence microscopy^29^. Similarly, the colocalization of Zo1 and F-actin has been observed in cultured cells using a 100× objective^47^. Spinning disk confocal super-resolution microscopy showed the co-occurrence of Zo1 and F-actin in the top half of mouse tracheal cells rather than the apical plane^48^. The zigzag pattern of Zo1 can be clearly visualized in the human corneal endothelium, while the F-actin signal appears blurred^49^ under a 100× objective in confocal microscopy. These findings establish the connection between Zo1 and F-actin, although image resolution needs improvement.^29^ In contrast, with the use of ColExM, our study successfully demonstrated the partial colocalization of Zo1 and F-actin in the corneal endothelium **(Fig. 4 and Supplementary video 7)**, corresponding to the actin-binding site of 28 amino acids in Zo1^29^. In addition to investigating tight junctions, ColExM can also be used in the future to decipher the underlying mechanisms of the basal pole of the corneal endothelium interdigitating and attaching to Descemet’s membrane.

Applications of ColExM collectively provide profound insights into the nanostructures of organelles, such as mitochondria and lysosomes, in collagen-abundant tissues under physiological and pathological conditions. In addition, ColExM with increasing expansion factors up to 10×^50^ may be used to clarify spatial sub-nano structures or molecular interactions, such as transcriptional factor translocation, for signal transduction in intact tissues.

## Methods

### Reagents and reagent preparation

Monomer solution and gelling solution were prepared as described by Asano et al.^2^ Detailed information on solutions used for expansion microscopy is provided in **Table 1**.

| Anchoring solution | Stock conc. | Amount | Final conc. | Supplier | Catalogue # |
| --- | --- | --- | --- | --- | --- |
| Methacrylic Acid *N*-Hydroxysuccinimide Ester (MA-NHS) | 10 mg/mL in DMSO |  | 0.1 mg/mL in 1 x PBS | Sigma | 730300 |
| Monomer solution | Stock conc. | Amount | Final conc. | Supplier | Catalogue # |
| Sodium Acrylate | 38% (w/w) | 2.25 mL | 8.6% (w/w) | Sigma | 408220 |
| Acrylamide solution | 40% (w/w) | 625 µL | 2.5% (w/w) | Sigma | A4058 |
| *N,N*-Methylenebisacrylamide solution (BIS) | 2% (w/w) | 750 µL | 0.15% (w/w) | Sigma | M1533 |
| Sodium Chloride | 5 M | 4 mL | 2 M | Sigma | 31434 |
| PBS | 10 x | 1 mL | 1 x |  |  |
| Double distilled water |  | 775 µL |  |  |  |
| Gelling solution | Stock conc. | Amount | Final conc. | Supplier | Catalogue # |
| Monomer solution |  | 940 µL |  |  |  |
| 4-Hydroxy-TEMPO (4HT) | 0.5% (w/w) | 20 µL | 0.01% (w/w) | Sigma | 176141 |
| Tetramethylethylenediamine (TEMED) | 10% (w/w) | 20 µL | 0.2% (w/w) | Sigma | T7024 |
| Ammonium Persulfate (APS) | 10% (w/w) | 20 µL | 0.2% (w/w) | Sigma | A3678 |
| Collagenase digestion buffer | Stock conc. | Amount | Final conc. | Supplier | Catalogue # |
| Collagenase II | 10 mg/mL | 125 µL | 0.25 mg/mL | Sigma | C6885 |
| Dispase II | 10 mg/mL | 500 µL | 1 mg/mL | Thermo Fisher | 17105041 |
| Hyaluronidase | 12.5 mg/mL | 500 µL | 1.25 mg/mL | Sigma | H3506 |
| Liberase | 25 mg/mL | 50 µL | 0.25 mg/mL | Sigma | 05401020001 |
| DNase I | 10 U/µL | 23.2 µL | 0.0464 U/µL | Sigma | 4716728001 |
| D-Sorbitol | 2.5 M | 200 µL | 0.1 M | Sigma | S1876 |
| HEPES | 10 mM | 25 µL | 0.05 mM | Sigma | H3375 |
| DMEM: F12 medium |  | 3.575 mL |  | Thermo Fisher | 21041025 |
| Digestion buffer | Stock conc. | Amount | Final conc. | Supplier | Catalogue # |
| Guanidine Hydrochloride | 8 M | 5 mL | 0.8 M | Sigma | G9284 |
| Triton X-100 | 100% | 250 µL | 0.5% (v/v) | Sigma | T9284 |
| Tris, pH 8.0 | 1 M | 2.5 mL | 50 mM | Thermo Fisher | AM9855G |
| Calcium Chloride Hexahydrate | 0.2 M | 500 µL | 2 mM | Sigma | 442909 |
| Double distilled water |  | 41.75 mL |  |  |  |
| Proteinase K digestion buffer | Stock conc. | Amount | Final conc. | Supplier | Catalogue # |
| Digestion buffer |  | 990 µL |  |  |  |
| Proteinase K | 800 U/mL | 10 µL | 8 U/mL | New England Biolabs | P8017S |

| 10 x PBS | Stock conc. | Amount | Final conc. | Supplier | Catalogue # |
| --- | --- | --- | --- | --- | --- |
| Sodium Chloride |  | 80 g | 8% (w/w) | Sigma | 31434 |
| Potassium Chloride |  | 2 g | 0.2% (w/w) | Sigma | 31248 |
| Sodium Phosphate Dibasic |  | 14.4 g | 1.44% (w/w) | Sigma | S0751 |
| Potassium Phosphate Monobasic |  | 2.4 g | 0.24% (w/w) | Sigma | P5379 |
| Double distilled water |  | 1 L |  |  |  |
| Fixation solution | Stock conc. | Amount | Final conc. | Supplier | Catalogue # |
| Paraformaldehyde (PFA) |  | 2 g | 4% (w/w) | Sigma | 158127 |
| 1 x PBS |  | 50 mL |  |  |  |
| Blocking buffer | Stock conc. | Amount | Final conc. | Supplier | Catalogue # |
| Normal Goat Serum (NGS) | 100% | 1.5 mL | 10% (v/v) | Sigma | NS02L |
| Sodium Azide (NaN<sub>3</sub>) | 1% (w/w) | 300 µL | 0.02% (w/w) | Sigma | S2002 |
| Triton X-100 | 20% (v/v) | 1.5 mL | 2% (v/v) | Sigma | T9284 |
| 1 x PBS |  | 11.7 mL |  |  |  |
| Dilution buffer | Stock conc. | Amount | Final conc. | Supplier | Catalogue # |
| Normal Goat Serum (NGS) | 100% | 500 µL | 1% (v/v) | Sigma | NS02L |
| Sodium Azide (NaN<sub>3</sub>) | 1% (w/w) | 1 mL | 0.02% (w/w) | Sigma | S2002 |
| Triton X-100 | 20% (v/v) | 625 µL | 0.25% (v/v) | Sigma | T9284 |
| 1 x PBS |  | 47.875 mL |  |  |  |
| Rinsing buffer | Stock conc. | Amount | Final conc. | Supplier | Catalogue # |
| Sodium Chloride |  | 30 g | 3% (w/w) | Sigma | 31434 |
| Triton X-100 | 20% (v/v) | 50 mL | 1% (v/v) | Sigma | T9284 |
| 1 x PBS |  | 950 mL |  |  |  |

### Mice

Eight-week-old C57BL/6 mice were purchased from the Taiwan National Laboratory Animal Center (NLAC), Taiwan. Thy1-YFP-H (B6.Cg-Tg(Thy1-YFP)HJrs/J, #003782)^51^ and R26R-Mito-EGFP (CDB0251K)^25^ mice were obtained from The Jackson Laboratory and Riken, respectively. Thy1-YFP-H mice express yellow fluorescent protein at high levels in motor and sensory neurons and subsets of central neurons. All procedures with mice were approved by the Institutional Animal Care and Use Committees of National Taiwan University (IACUC Approval No: NTU-111-EL-00112).

### Corneal and skin sample preparation

C57BL/6, Thy1-YFP-H, and Mito-EGFP mice were sacrificed to collect corneal and skin specimens for expansion microscopy. Hair shafts were removed using a hair removal cream (Veet) to reduce melanin interference. Skin samples were collected from the dorsal skin and footpads. The corneal and skin samples collected were further fixed using 4% paraformaldehyde (PFA) in phosphate-buffered saline (PBS; Santa Cruz, sc-281692) at 4 °C for 24 hours. The corneas were dissected carefully under a dissecting microscope and the intraocular lens, iris, retina, and conjunctiva were discarded. The PFA-fixed tissues were washed with PBS three times and then stored in PBS at 4 °C before expansion.

### Ex vivo corneal organ culture for disrupting tight junctions in the corneal endothelium

To disrupt tight junctions, corneas were dissected from enucleated eyeballs while preserving the conjunctiva to avoid injury to the corneal endothelium. Corneas were incubated in Dulbecco’s Modified Eagle Medium (Thermo Fisher, 21063029) with 2 mM EGTA (Sigma, E3889) at room temperature for 30 minutes, 1 hour, or 2 hours. After treatment, corneas were washed with PBS and fixed with 4% PFA at room temperature for 15 minutes. The conjunctiva and irises were dissected under a dissecting microscope after fixation. The samples were then bathed in PBS at 4 °C until performance of ExM.

### Immunofluorescence staining

Tissue sections (∼400 μm in thickness) were first incubated in blocking buffer at 4 °C overnight before incubation in primary antibodies in dilution buffer at 4 °C for 48 hours. The sections were then incubated with secondary antibodies, which were conjugated with fluorescent proteins or biotin, in dilution buffer at 4 °C for 24 hours. Biotinylated samples were further incubated with streptavidin-conjugated fluorescent dyes at 4 °C for 24 hours. The samples were rinsed by incubating them in rinsing buffer at room temperature (20 minutes × 3) after each antibody incubation. The sections were immersed in 4% PFA for post-fixation at room temperature for 15 minutes at the end of immunostaining. Nuclear counter-staining was performed with DAPI (1:500 in double-distilled water; Sigma, D9542) or propidium iodine (1:500 in double-distilled water; Biotium, 40017) at 4 °C overnight. Stained tissues were then washed twice for 5 minutes each with PBS to remove excess DAPI or propidium iodine. Detailed information on solutions for immunofluorescence staining is provided in **Table 1** and **Supplementary Table 1.**

### 4.5× tissue expansion of collagen-abundant tissues

All stained tissues were immersed in methacrylic acid *N*-hydroxysuccinimide ester at 4 °C overnight. Then, before gelling, anchored tissues were incubated in monomer solution twice at 4 °C for 5 minutes each, followed by two other incubations (for 5 minutes and then 25 minutes) in the gelling solution at 4 °C. The tissues were later transferred to a custom-made gelation chamber (5–8 layers of reinforced O-rings stacked to make a 370–590 μm thick spacer on the glass slide) containing 30–35 μL gelling solution. Next, the chamber was covered with glass coverslips and placed in a humidified container at 37 °C for 2 hours. Each gel was then incubated in 1 mL of collagenase digestion buffer at 37 °C for 48 hours (with the digestion buffer replaced with fresh buffer every 24 hours), followed by incubation in proteinase K digestion buffer at 55 °C for another 48 hours (with the buffer replaced with fresh buffer every 24 hours). The gels were then washed three times for 15 minutes each with double-distilled water at room temperature with shaking to fully expand the gel (∼4.5× expansion). The workflow for ColExM with protein retention is shown in **Supplementary Figure 5**.

### Volumetric imaging and 3D visualization

Unexpanded samples were immersed in NFC2 solutions (Nebulum, Taipei, Taiwan) to match the refractive index. Fully expanded gels were immersed in double-distilled water for imaging. Tissue slices were imaged using a laser-scanning microscope (ZEISS LSM 880 with Airyscan, Germany) and a 40×/1.0 NA water immersion lens before expansion (**Fig. 1, e and g**). Otherwise, all other images were acquired with a spinning disk confocal microscope (Andor Dragonfly 200, Oxford Instruments, UK) equipped with a motorized stage, 405/488/561/637nm laser lines, and 5×/0.1NA air and 10×/0.3NA and 40×/0.8NA water immersion lenses. Pre-ExM and post-ExM images of tight junctions were acquired with the 40× water immersion lens from different corneal samples to avoid severe photobleaching during image acquisition (**Figs. 5 and 6 and Supplementary Fig. 4**). After image acquisition, Imaris software (Imaris 9.6.0, Bitplane, Belfast, UK) was employed for image processing and 3D visualization.

### Super-resolution image acquisition and visualization

We acquired super-resolution images of tight junctions using a stimulated emission depletion microscope (STED) (Leica TCS SP8 STED, Germany). Immunostaining of corneas was performed according to standard procedure.^52^ The corneas were mounted in Dako mounting medium (Dako). We replaced the Alexa Fluor 488 dye-conjugated antibodies used in confocal microscopy with STED-dedicated secondary antibodies (abberior STAR RED, abberior STAR 580, Abberior STAR 635, and phalloidin) to avoid photobleaching caused by the depletion laser. During image acquisition, the excitation and emission wavelengths were set according to the recommendations from the manufacturer. In addition, we used a 750 nm depletion laser with 10% laser power to perform depletion of three secondary antibodies. We used an abberior NANOPARTICLE SET to calibrate the system prior to imaging for high-resolution performance. After image acquisition, Imaris software (Imaris 9.6.0, Bitplane, Belfast, UK) was employed for image processing and 3D visualization.

### Measurement of root-mean-square error

Quantitative distortion analysis was performed as described by Chozinski, et al.^3^ The same fields of view were imaged before and after expansion. Post-ExM images were initially aligned with their corresponding pre-ExM images through rotation, translation, and uniform scaling using *Imaris* and *Fiji*. Subsequently, *Register Virtual Stack Slices* was applied to post-ExM images for matching with pre-ExM images. The registered images were further processed using custom-written Mathematica scripts.

## Acknowledgments

We thank the Brain Research Center, National Tsing-Hua University, Taiwan, for the spinning disk confocal and Airyscan technical assistance. In addition, we thank the imaging core at the First Core Lab, National Taiwan University College of Medicine, for technical support in STED image acquisition. This work was supported by Taiwan Bio-Development Foundation (TBF) (to SJL), National Taiwan University (111L892401 to SJL), National Taiwan University Hospital (111IF0006 to SJL), Taiwan National Health Research Institutes (NHRI-EX111-11112EI to SJL) and National Science and Technology Council (NSTC-111-2636-B-007-007, NSTC-112-2326-B-007-006 to LAC; 111-2327-B-002-015 and 110-2314-B-002-190-MY3 to SJL). This work was also supported by the Brain Research Center under the Higher Education Sprout Project, co-funded by the Taiwan Ministry of Education and Taiwan National Science and Technology Council. YFW was supported by a postdoctoral fellowship from Taiwan National Science and Technology Council (111-2811-B-002-156 and 112-2811-B-002-063).

## Author information

## Contributions

Conceptualization: SJL, LAC

ColExM experiment: YHC

EGTA drug treatment on cornea: YFW and YXZ

Confocal image preparation: YHC

STED image preparation: SCH

Image processing and analysis: YHC

Animal dissection and sample preparation: YFW and YHL

Writing: YHC, YFW, SJL and LAC

Reviewing and editing: YHC, YFW, SJL and LAC

Supervision: SJL, LAC

## Corresponding authors

Correspondence to SJL and LAC

