## Supplementary Fig.1-5, Supplementary Table 1 for "Expansion microscopy for super-resolution imaging of collagen-abundant tissues"

&

Li-An Chu, PhD
Department of Biomedical Engineering and Environmental Sciences, National Tsing Hua University
101, Section 2, Kuang-Fu Road, Hsinchu, Taiwan 300
Tel: (+886)-3-5725077 ext. 42681 Fax: (+886)-3-5718649

Orcid: 0000-0002-4092-3024

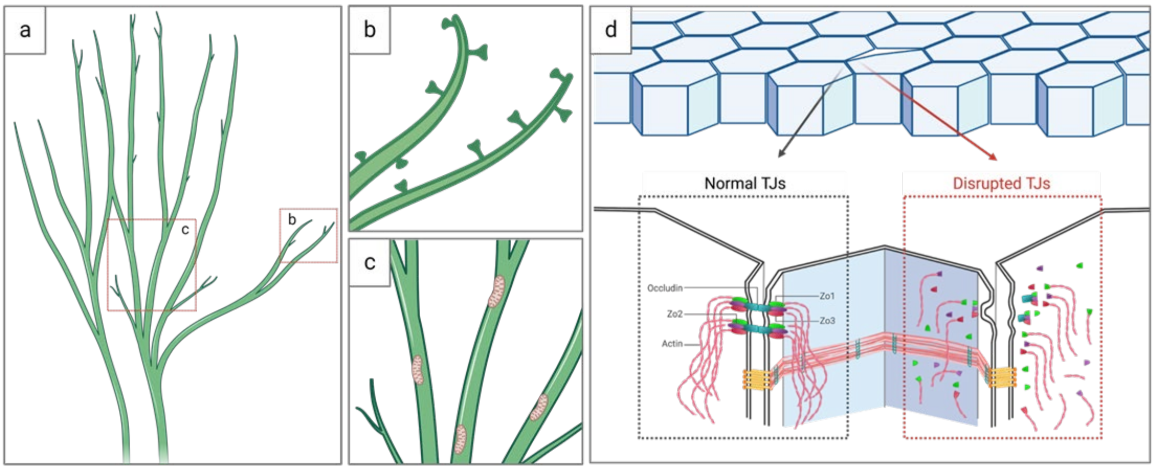


**Graph summary**

ColExM unveils nanostructures in the cornea and skin. It enables 3D visualization of nerves innervating the cornea and hair follicles (a), revealing spine-like structures (b) and mitochondria (c) along the nerves. Furthermore, it allows for the observation of conformational changes in the nanoscale structure of the intracellular domain of tight junctions within the corneal endothelium following EGTA treatment (d).

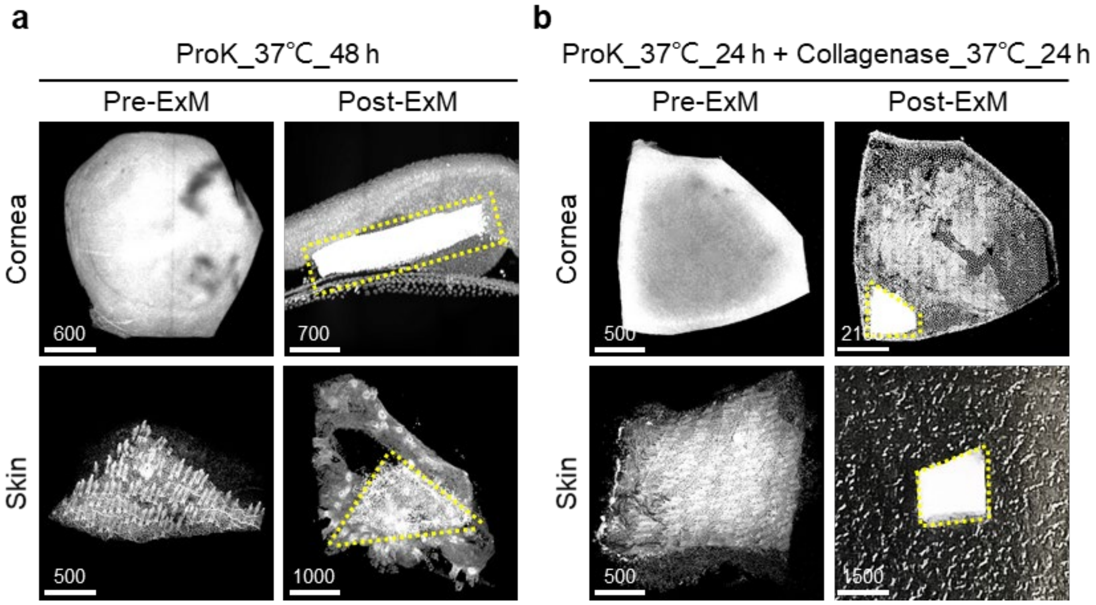


**Supplementary Fig. 1 | Collagen-abundant tissue expansion based on previous ExM protocols.** **a-b,** Representative images of pre- and post-expansion in cornea and skin. Nuclear PI staining was shown in gray. Non-expanded structures were marked with yellow dotted lines. Scale bar unit: µm. Scale bars are not adjusted for tissue expansion.

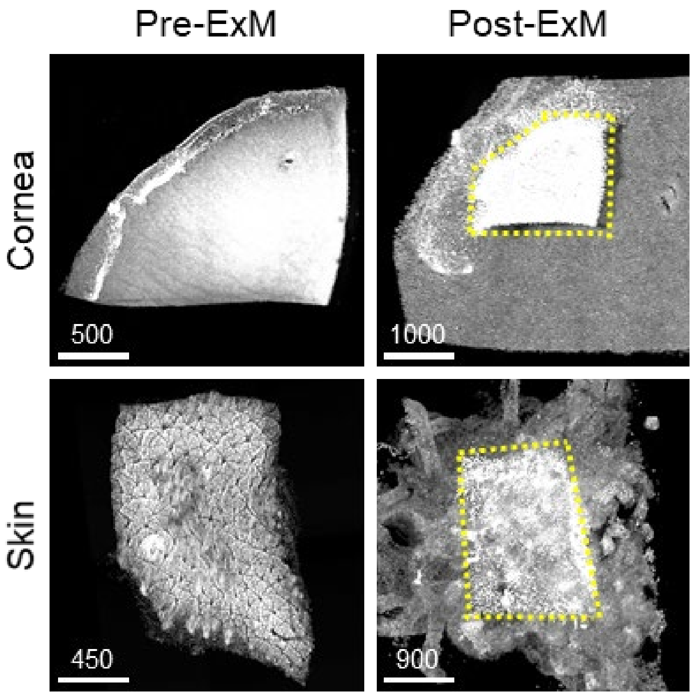


**Supplementary Fig. 2** | Collagen-abundant tissue expansion under pre-treatment of 15-minute collagenase before fixation. Representative images of pre- and post-expanded cornea and skin. Nuclear PI staining was shown in gray. Non-expanded structures were marked with yellow dotted lines. Scale bar unit: µm. Scale bars are not adjusted for tissue expansion.

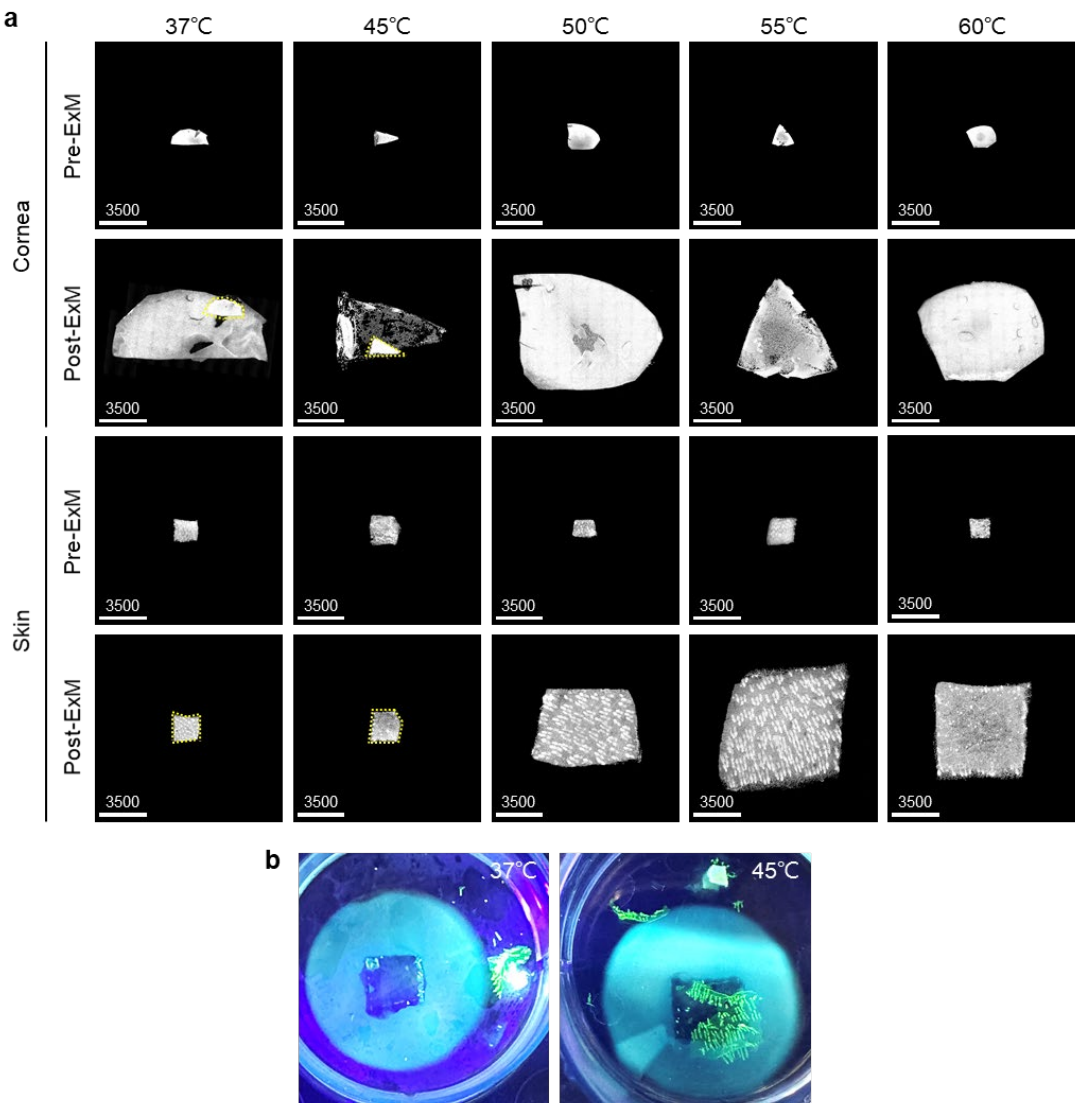


**Supplementary Fig. 3** | Collagen-abundant tissues were expanded by a 48-hour collagenase digestion at 37°C, followed by a 24-hour ProK digestion under various temperature conditions **a,** Representative images of pre- and post-expanded cornea and skin processed at 37°C, 45°C, 50°C, 55°C and 60°C for 24-hour ProK digestion. Nuclear SYTO16 staining was shown in gray. Non-expanded structures were marked with yellow dotted lines. Scale bar unit: μm. **b,** Representative images of skin were obtained following a 48-hour collagenase digestion at 37°C, followed by a 24-hour ProK digestion at 37°C (left panel) and 45°C (right panel).

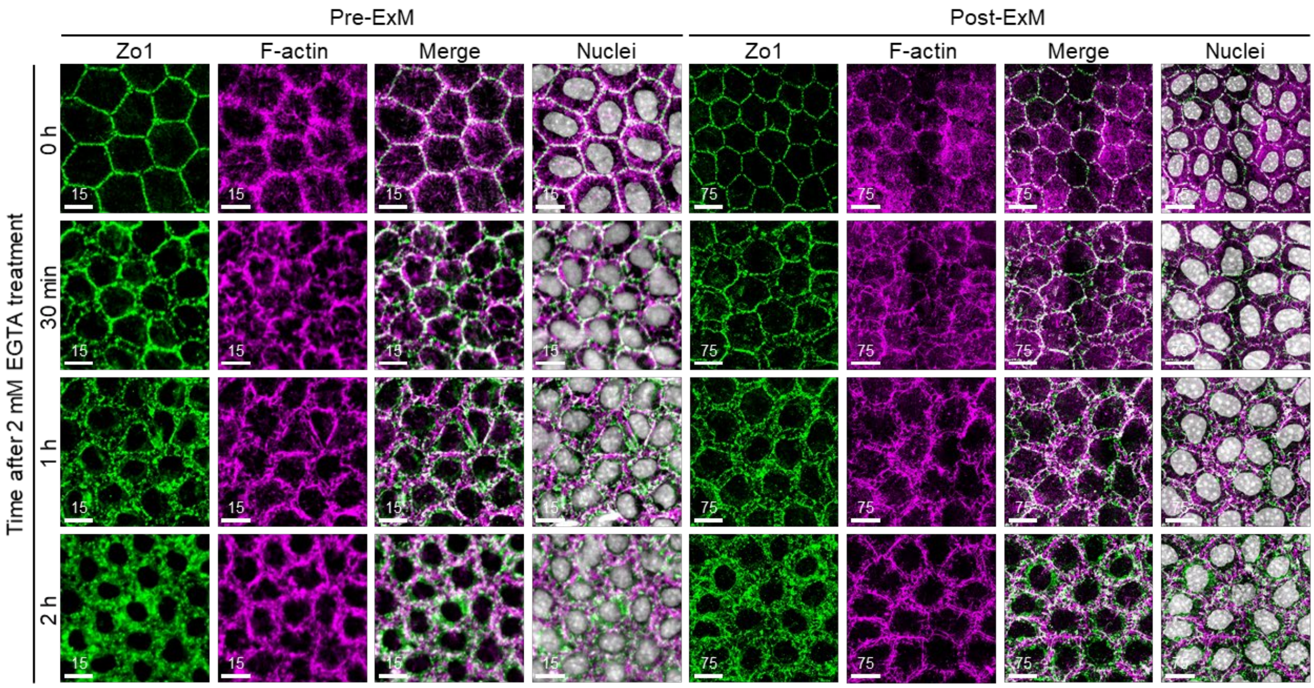


**Supplementary Fig. 4 | Tight junction of corneal endothelial cells conformationally changes after disruption by EGTA.** Expression of Zo1 (green), F-actin (magenta) and nuclei (gray) in corneal endothelium before and after expansion following EGTA treatment. Colocalization of Zo1 and F-actin was shown in white in merge images. Scale bar unit: µm. Scale bars have not been adjusted for expansion factor.

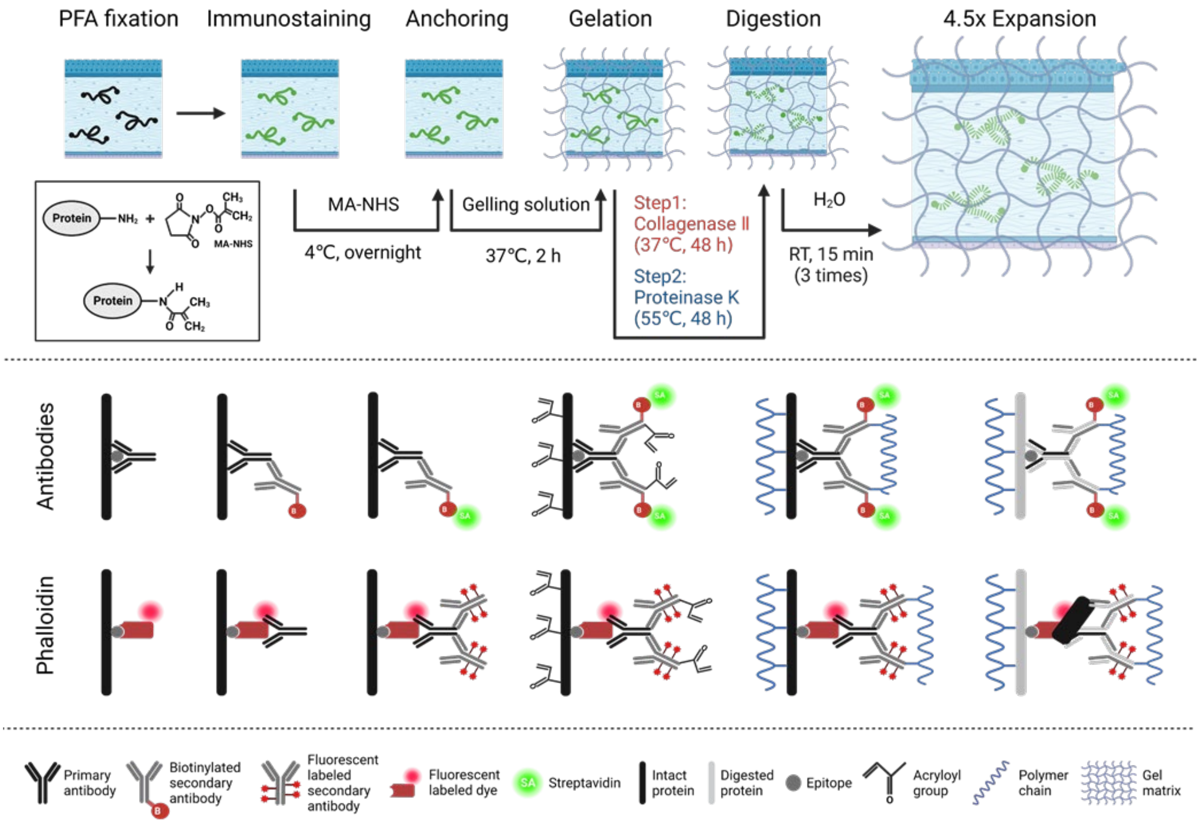


**Supplementary Fig. 5 | The workflow for ColExM with protein retention.** Collagen-abundant mouse tissues are initially fixed by PFA and are immunostained to target specific structures. Methacrylic acid *N*-hydroxysuccinimide ester (MA-NHS) is used to anchor proteins of the entire sample and fluorophore-labeled antibodies to polymer-linking groups of the hydrogel during gelation. The samples then undergo sequential homogenization with collagenase and ProK digestion buffers, followed by incubation in double distilled water, resulting in approximately 4.5x expansion in three dimensions.

| Primary Antibody | Host | Supplier | Catalogue # | Dilution | Incubation Time |
|---|---|---|---|---|---|
| Zo1 | Mouse | Thermo Fisher | 33-9100 | 1:200 | 48 + hrs at 4℃ |
| GFP | Chicken | Abcam | ab13970 | 1:200 | 48 + hrs at 4℃ |
| Fluorescein/Oregon Green | Rabbit | Thermo Fisher | A-889 | 1:100 | 48 + hrs at 4℃ |
| Tubulin β3 | Rabbit | CST | 5568S | 1:100 | 48 + hrs at 4℃ |
| **Secondary Antibody** | **Host** | **Supplier** | **Catalogue #** | **Dilution** | **Incubation Time** |
| Goat-anti-Mouse-Biotin-SP | Goat | Jackson Lab | 715-067-033 | 1:200 | 24 + hrs at 4℃ |
| Goat-anti-Chicken-Biotin-SP | Goat | Jackson Lab | 103-065-155 | 1:200 | 24 + hrs at 4℃ |
| Goat-anti-Rabbit-Biotin-SP | Goat | Jackson Lab | 111-065-003 | 1:200 | 24 + hrs at 4℃ |
| Goat-anti-Chicken-Alexa-488 | Goat | Jackson Lab | 103-545-155 | 1:200 | 24 + hrs at 4℃ |
| Goat-anti-Rabbit-Alexa-633 | Goat | Thermo Fisher | A-21071 | 1:200 | 24 + hrs at 4℃ |
| **Staining Reagent** | | **Supplier** | **Catalogue #** | **Dilution** | **Incubation Time** |
| Fluorescein phalloidin | | Thermo Fisher | F432 | 1:40 | 120 + hrs at 4℃ |
| SA-Alexa 488 | | Jackson Lab | 016-540-084 | 1:200 | 24 + hrs at 4℃ |
| SA-Rhodamine X red | | Jackson Lab | 016-290-084 | 1:200 | 24 + hrs at 4℃ |
| SYTO™16 | | Thermo Fisher | S7578 | 1:500 | 18 + hrs at 4℃ |
| Propidium Iodine (PI) | | Biotium | 40017 | 1:500 | 18 + hrs at 4℃ |
| DAPI | | Sigma | D9542 | 1:500 | 18 + hrs at 4℃ |

**Supplementary Table 1**

Antibodies and conditions. Table showing primary and secondary antibody, staining reagent conditions used for expansion microscopy.
